# Bacterial single-cell genomics enables phylogenetic analysis and reveals population structures from *in vitro* evolutionary studies

**DOI:** 10.1101/2020.08.25.266213

**Authors:** Matt Bawn, Johana Hernandez, Eleftheria Trampari, Gaetan Thilliez, Mark A. Webber, Robert A. Kingsley, Neil Hall, Iain C. Macaulay

## Abstract

Single-cell DNA sequencing has the potential to reveal detailed hierarchical structures in evolving populations of cells. Single cell approaches are increasingly used to study clonal evolution in human ageing and cancer, but have not yet been deployed to study evolving microbial populations. Here, we present an approach for single bacterial genomic analysis using FACS isolation of individual bacteria followed by whole-genome amplification and sequencing. We apply this to *in vitro* experimental evolution of a hypermutator strain of *Salmonella* in response to antibiotic stress (ciprofloxacin). By analysing sequence polymorphisms in individual cells from the population we identified the presence and prevalence of sub-populations which have acquired polymorphisms in genes previously demonstrated to be associated with ciprofloxacin susceptibility. We were also able to identify that the population exposed to antibiotic stress was able to both develop resistance whilst maintaining diversity. This population structure could not be resolved from bulk sequence data, and our results show how high-throughput single-cell sequencing can enhance experimental studies of bacterial evolution.

## Main Text

A long-term evolution experiment in *E. coli* has emphasised the insights that can be gained from the study of evolutionary processes in bacteria *in vitro* (Tenaillon et al., 2016). Experimental evolution studies have the power to transform our understanding of the molecular mechanisms underpinning the emergence of phenotypic traits including resistance to antimicrobial compounds (Remigi et al., 2019). These experiments are typically analysed by either characterising individual isolates, or by assessing the genetic changes occurring in populations through the bulk sequencing of cultured samples. Neither approach can fully resolve the genetic heterogeneity present within bacterial communities; to do so requires single-cell genomic analysis with sufficient coverage and resolution to capture single-nucleotide polymorphisms (SNPs).

Single-cell studies on bacterial systems highlighted the applicability of Multiple Displacement Amplification (MDA, (Dean et al., 2002)) to provide enough material for genome analysis (Raghunathan et al., 2005) and the potential to sequence uncultured isolated cells (Marcy et al., 2007). Subsequent studies focused on *de novo* genome assembly from single cell sequencing (Rodrigue et al., 2009; Woyke et al., 2010) and technology development to facilitate this (Gole et al., 2013). Nevertheless, the majority of single-cell genomic technology development to date has involved eukaryotic systems, and high-throughput effective solutions dedicated to microbial systems remain a largely unmet need (Mincarelli et al., 2018), and the application of single cell sequencing to resolve population structure in an evolving community has yet to be explored.Here, we present an approach to investigate the population structure of bacteria under selection for antimicrobial resistance by isolation and single-cell genome sequencing of hundreds of individual bacteria from planktonic cultures by Fluorescence Activated Cell Sorting (FACS) followed by MDA (**Figure 1A**).

**Figure 1:**
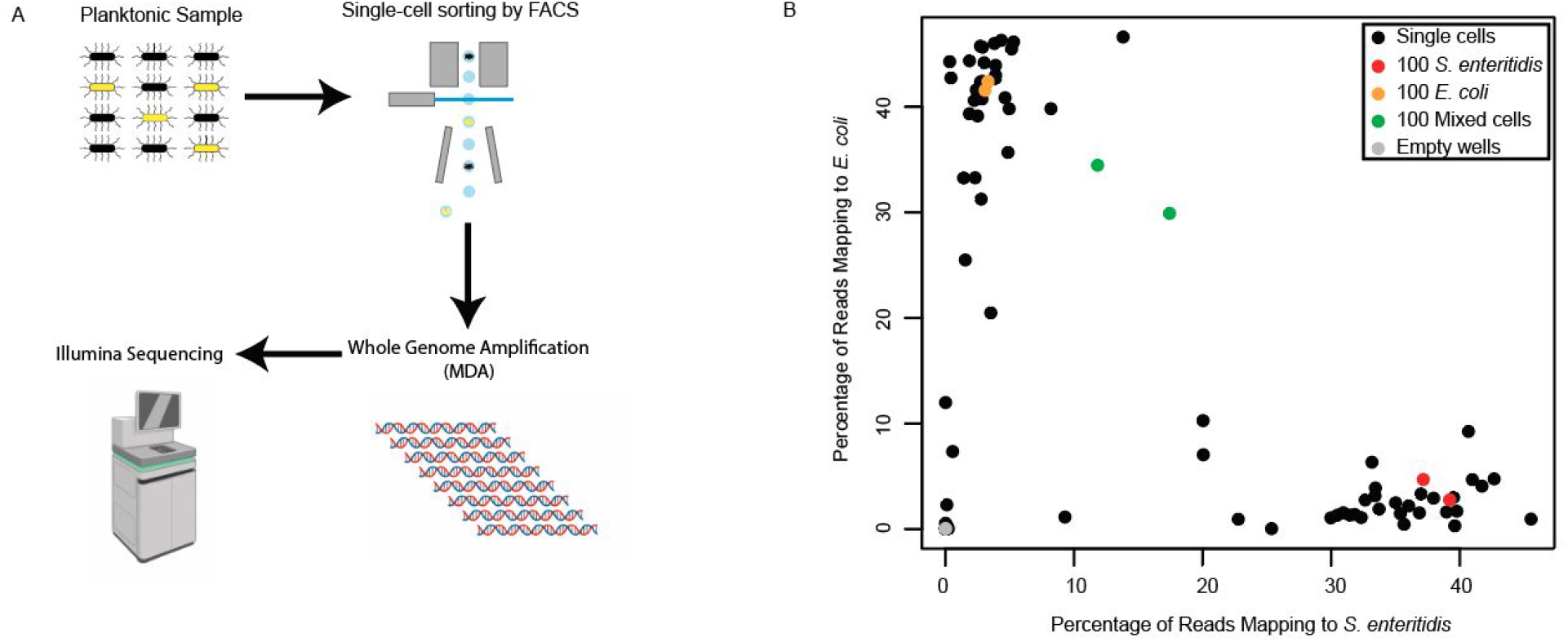
**A**) Overview of approach. Individual bacteria were sorted from planktonic samples by FACS, followed by MDA and Illumina sequencing B) Barnyard analysis in which single cells from a 1:1 mixture of *E. coli* and *S*. Enteritidis were isolated by FACS and sequenced. The percentage of reads mapping to each genome for each single cell (black), empty wells (grey) and multi-cell controls (red - *S*. Enteritidis, orange - *E. coli* and green - mixed), are shown.

To test the ability to sort individual bacteria, we performed a “barnyard” experiment, sorting single events from a 1:1 mixture of *E. coli* and *S. enterica* serotype Enteritidis (*S.* Enteritidis) into a 96 well plate, followed by whole genome amplification and sequencing. Individual cells predominantly had reads mapping to either one or the other genome, indicating that individual bacteria can be sorted from a mixture with a low rate of doublet cells (**Figure 1B**). Empty wells generated negligible amounts of mapping reads, while deliberately mixed mini-bulk wells showed a mixture of reads mapping to each genome.

To test the ability of this approach to detect genomic heterogeneity and study population structure, we generated test populations founded by a wild type (WT) *S.* Enteritidis, or its isogenic hypermutator *mutS* deletion mutant (*mutS*). Both strains were repeatedly passaged in the presence or absence of half of the Minimum Inhibitory Concentration (MIC) of ciprofloxacin twice a day over two weeks for ~150 generations. We chose this experimental model as the progression of genome variation at the SNP level associated with ciprofloxacin resistance has been well characterised (Redgrave et al., 2014). We expected to see a difference in diversity between drug-exposed and naive populations in these conditions with selection in the former potentially imposing a bottleneck on drug-sensitive lineages. Measurement of the MICs for the final populations revealed no change in ciprofloxacin susceptibility in untreated lineages but emergence of resistant mutants in all those populations exposed to the antibiotic (**Supplementary Table 1**). Isolation and susceptibility testing of randomly selected individual colonies from the final populations confirmed the emergence of resistant variants in the exposed lineages. This analysis also showed heterogeneity within the population with 13.6% of isolates being resistant (MIC >0.06 mg/L) to ciprofloxacin in the exposed lineage (**Supplementary Figure 1**).

We sorted and sequenced 352 single WT and *S.* Enteritidis *mutS* cells, from populations cultured in the presence or absence of ciprofloxacin (88 cells from each group), as well as mini-bulk (100 cells) and empty wells for each experimental group. In parallel, we performed high-coverage Polymerase Chain Reaction (PCR) free sequencing of the cultured populations to determine high-confidence variant calls (**Figure 2A**).

**Figure 2:**
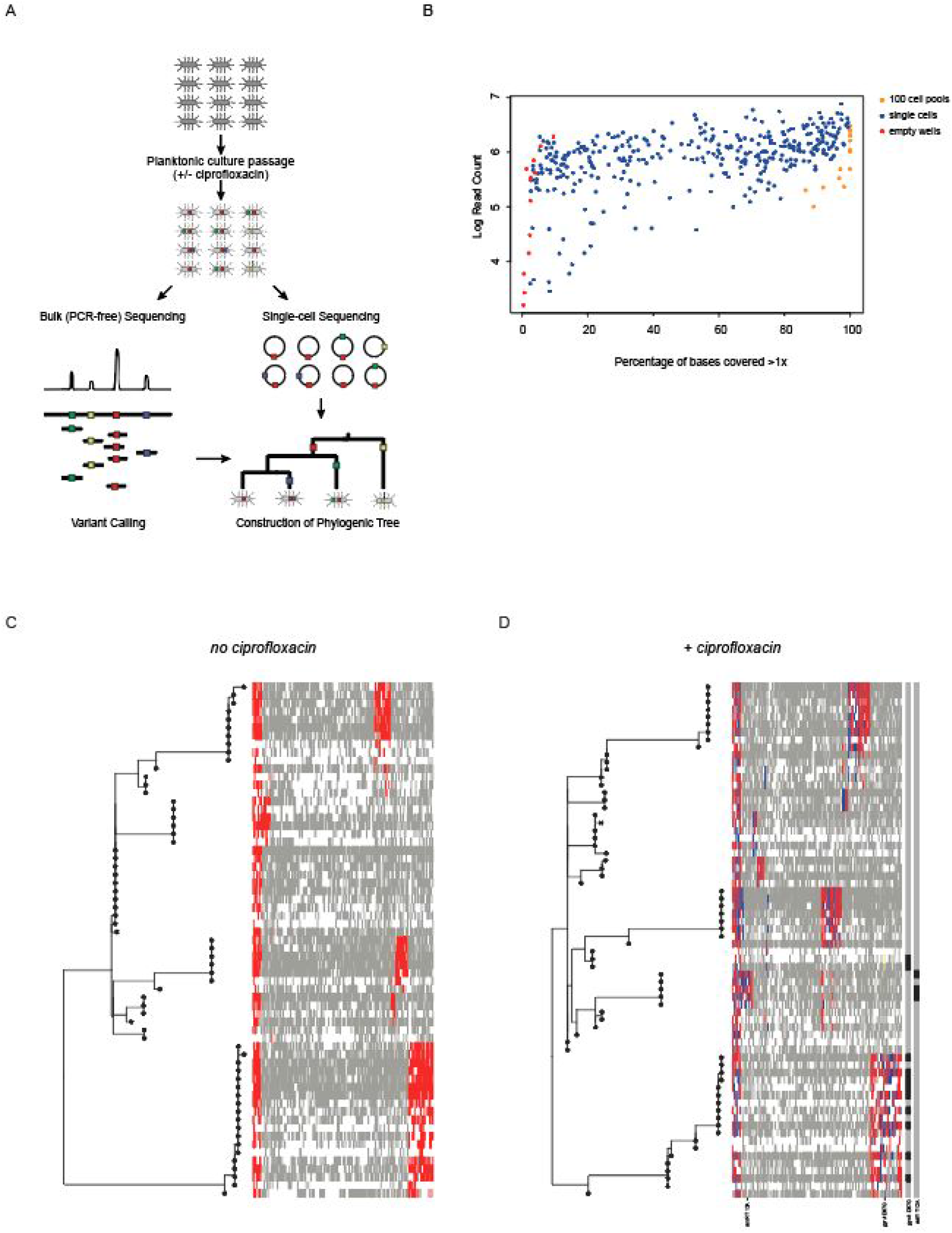
A) Experimental overview, *S.* Enteritidis was cultured in the absence and presence of sub-inhibitory concentrations of ciprofloxacin for around 100 generations. Bulk, PCR-free sequencing and single-cell sequencing of individual cells from the planktonic culture were then performed. B) Coverage plot for all whole genome amplified samples. C) and D) Phylogenies and population structure of hypermutator strains evolved in the (C) the absence and (D) presence of ciprofloxacin. The heatmap to the right of each phylogeny indicates the coverage and SNP status of each site in each sample used to construct the phylogeny (red: SNP, grey: coverage but no variant, white: no coverage). In the figure on the left, SNPs also present in the absence of ciprofloxacin are indicated in blue. The SNPs in *gyrA* and *acrR* are also highlighted.

In many of the single-cell samples, over 40% of bases of the *S.* Enteritidis genome were sequenced, and in some cases, near-complete genome coverage was obtained from single cells (**Figure 2B**). The mean percentage of reads that aligned to the *S.* Enteritidis reference genome for single-cells was 69%, with a mean coverage of 64X. PCR-free bulk libraries were sequenced to a depth of > 1000X. In the bulk sequencing data 2,753 SNPS were called at 1,500 unique genomic sites for the four samples, with more transitions seen in the *mutS* (mean proportion = 0.53) than in the W strain (mean proportion = 0.23) (**Supplementary Figure 2**). This pattern is expected due to the *mutS* deletion in the hypermutator and was previously seen in the *mutS* hypermutator strain of *S.* Enteritidis that evolved within an immunocompromised patient (Klemm et al., 2016).

In the corresponding single cell genome data, 115,286 raw SNPs were called at 99,617 unique genome sites. SNP profiles were similar for all four samples and distinct to those seen in bulk sequencing. Single-cell SNPs were dominated by G-A and C-T transitions (mean proportion 0.42) and C-A and G-T transversions (mean proportion 0.38) (**Supplementary Figure 2**). The prevalence of C-T transitions in MDA has been previously described and attributed to cytosine deamination during cell lysis (Chen et al., 2017) and G-T substitutions resulting from oxidation of guanine residues (Cheng et al., 1992).

We filtered the single-cell SNP data to remove as many false positives as possible. As true single-cell SNPs should be monoallelic, single cell SNPs present at frequencies below 0.9 were then removed (**Supplementary Figure 3**), comprising 105,684 SNPs (92% of initial single cell SNPs) at 93,239 positions, leaving 9,602 SNPs at 6,687 positions remaining. Inspection of the remaining SNP profiles showed that there was still an increased signal from G-A and C-T transitions when compared to the profile from the bulk sequencing −4,981 more G-A and C-T SNPs than A-G and T-C SNPs in the single-cell samples. Assuming the true number of transition types should be approximately the same as seen in the bulk data, we inferred that many of these additional SNPs were false positives. To remove potential false positives from further analysis, G-A and C-T SNPs seen in only one single-cell genome were removed, leaving 4,448 SNPs at 1,536 positions. Of these SNP positions, 234 (15%) also contained SNPs in the bulk sequence data. The mean frequency of bulk SNPs with positions also seen in the single cell SNPs is 0.18, compared with 0.03 for SNPs at positions not seen in single cells. Thus, SNPs observed by single-cell sequencing were in most cases not frequent enough to be observed by bulk sequencing, reflecting the limited sampling of each population (100s of cells) and the depth of sequencing of each cell.

The profiles of SNP transitions as well as their mean frequencies and the GC content of the SNP regions in single-cell only, high-coverage pooled and shared SNPs were also determined **Supplementary Figure 3**. To investigate the utility of the increased resolution that our method gives to population analyses we used a set of 316 high confidence SNPs in coding sequences of the single-cell genomes to elucidate phylogeny and population structure in the *mutS* strain in the absence and presence of ciprofloxacin (**Figure 2C and 2D**).

Since one of the populations was evolving in the presence of ciprofloxacin, we expected to observe SNPs in genes previously demonstrated to be associated with ciprofloxacin resistance. Examining the variants seen in the bulk sequencing data in the hypermutator strain (recovered from the population passaged with a 1:1000 dilution rate), SNPs were present in *gyrA* D87G (frequency 0.14), *acrR* T12A (frequency 0.06), *envC* G347S (frequency 0.014) and *ompX* A9V (frequency 0.011). In the corresponding single-cell data the *gyrA* D87G mutation was seen in 11 out of 88 (a frequency of 0.125) *mutS* cells grown in the presence of ciprofloxacin (**Figure 2D**). These frequencies are in agreement with those seen in the bulk sequencing data as well as the frequency of resistant isolates experimentally observed in the populations and show that the emergence and prevalence of resistant mutants was accurately detected by the single-cell approach (**Supplementary Figure 1**). The *acrR* T12A SNP was seen in three single-cell genomes (a frequency of 0.034) (**Figure 2D**). The *envC* and *ompX* mutations seen in the pooled data at minor frequencies (~ 0.01) were not observed in the single-cell genomes, likely due to their low frequency.

The population structure analysis of the hypermutator strain determined six second-level clades were present in the cultures without ciprofloxacin (**Figure 2C**) and nine in the presence of the antibiotic (**Figure 2D**), consistent with an unexpected increase in diversity under selection. This greater diversity in the drug-exposed culture suggested that a population exposed to a sub-inhibitory concentration of antibiotic can both develop resistance whilst maintaining overall diversity, since under these conditions no significant bottleneck was imposed on the population. This highlighted the power of single-cell approaches to dissect complexity in cell populations that would not be visible if only bulk sequencing had been performed, in part due to the inability to co-locate SNPs within individual genomes. Several SNPs were present in populations following culture with or without supplementation with ciprofloxacin that are likely due to adaptation to the media.However, some were unique to the *mutS* strain grown with antibiotic, and sub-clones of *gyrA* D87G and *acrR* T12A mutant cells were present as distinct clades on the phylogenetic tree (**Figure 2D**). The phylogenetic analysis provided an evolutionary roadmap for the bacterial populations and the framework for the imputation of variants in incompletely covered genomes.

In summary, we have demonstrated that FACS-based single-cell isolation and whole genome amplification of bacterial genomes can be readily performed on hundreds of cells derived from complex planktonic cultures. This approach enables the high-throughput and reliable determination of SNPs from individual bacteria within a population, which when combined with *in vitro* evolution experiments can be used to resolve population structure in evolving populations. This method allows greater understanding of population dynamics and will help predict evolutionary trajectories for populations exposed to stresses of interest.

---

**Supplementary Table 1:**
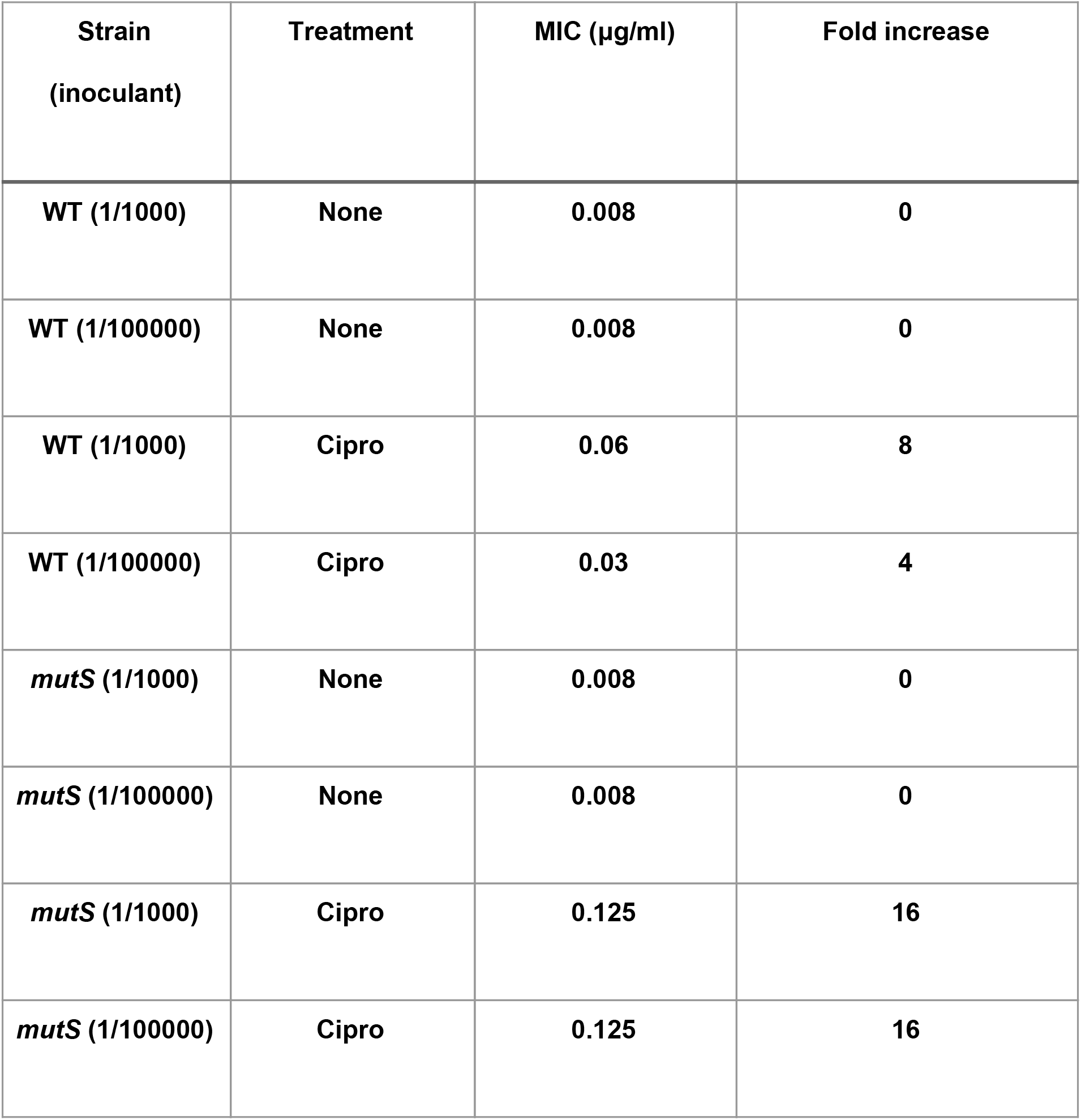
Details of strains and culture conditions used in these experiments.

| **Strain**<br>**(inoculant)** | **Treatment** | **MIC (µg/ml)** | **Fold increase** |
| --- | --- | --- | --- |
| **WT (1/1000)** | **None** | **0.008** | **0** |
| **WT (1/100000)** | **None** | **0.008** | **0** |
| **WT (1/1000)** | **Cipro** | **0.06** | **8** |
| **WT (1/100000)** | **Cipro** | **0.03** | **4** |
| ***mutS* (1/1000)** | **None** | **0.008** | **0** |
| ***mutS* (1/100000)** | **None** | **0.008** | **0** |
| ***mutS* (1/1000)** | **Cipro** | **0.125** | **16** |
| ***mutS* (1/100000)** | **Cipro** | **0.125** | **16** |

**Supplementary Figure 1.**
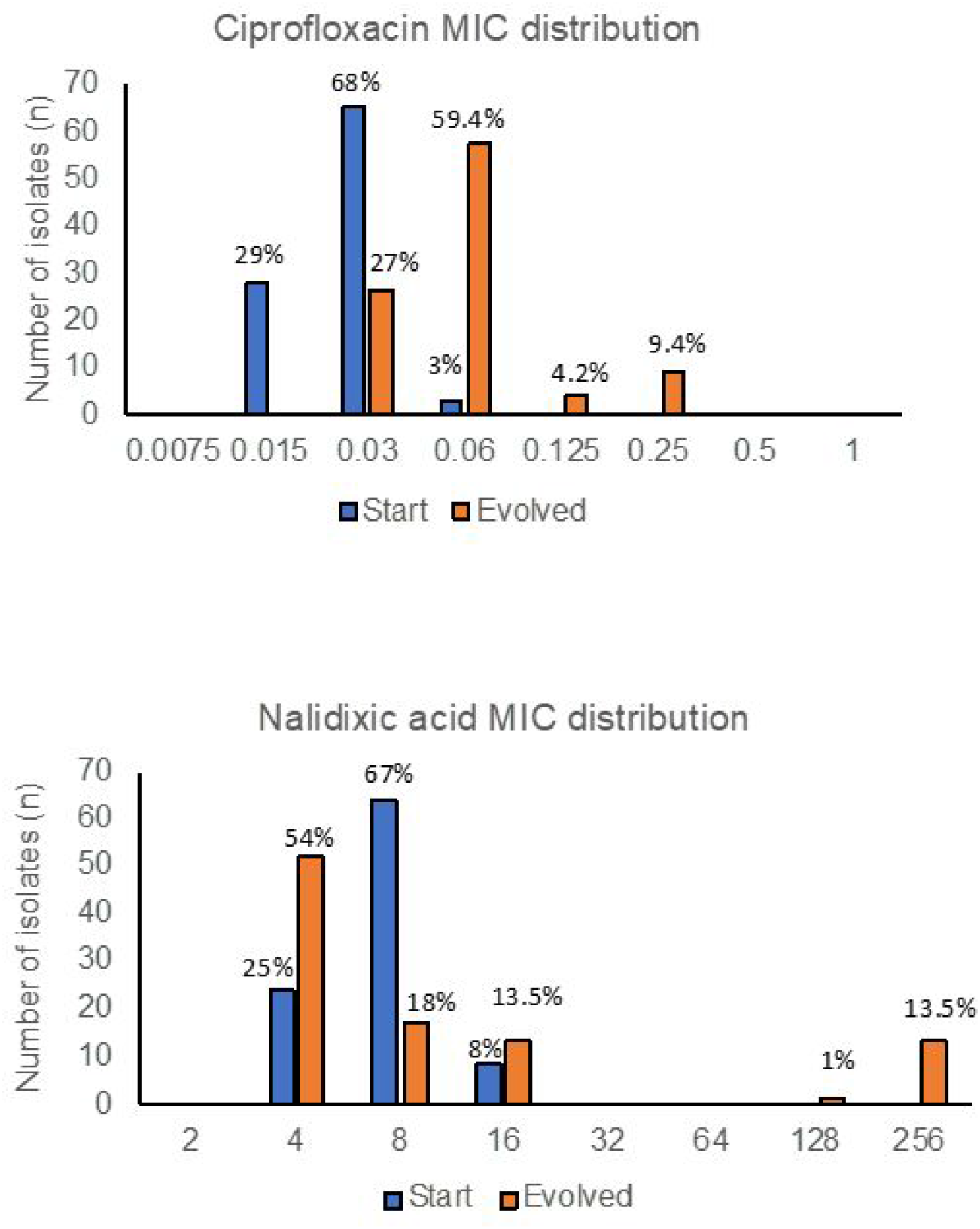
Heterogeneity of susceptibility of isolates to ciprofloxacin within populations. 96 individual colonies were isolated from *mutS* lineages (exposed and control) and the MIC of ciprofloxacin (top) and nalidixic acid (bottom) against all isolated strains was determined to identify the prevalence of resistance within the population. This confirmed selection of resistance in the exposed lineage, and identified heterogeneity within the population with 13.6% of individual isolates being resistant (MIC >0.06 mg/L) to ciprofloxacin in the exposed lineage.

**Supplementary Figure 2.**
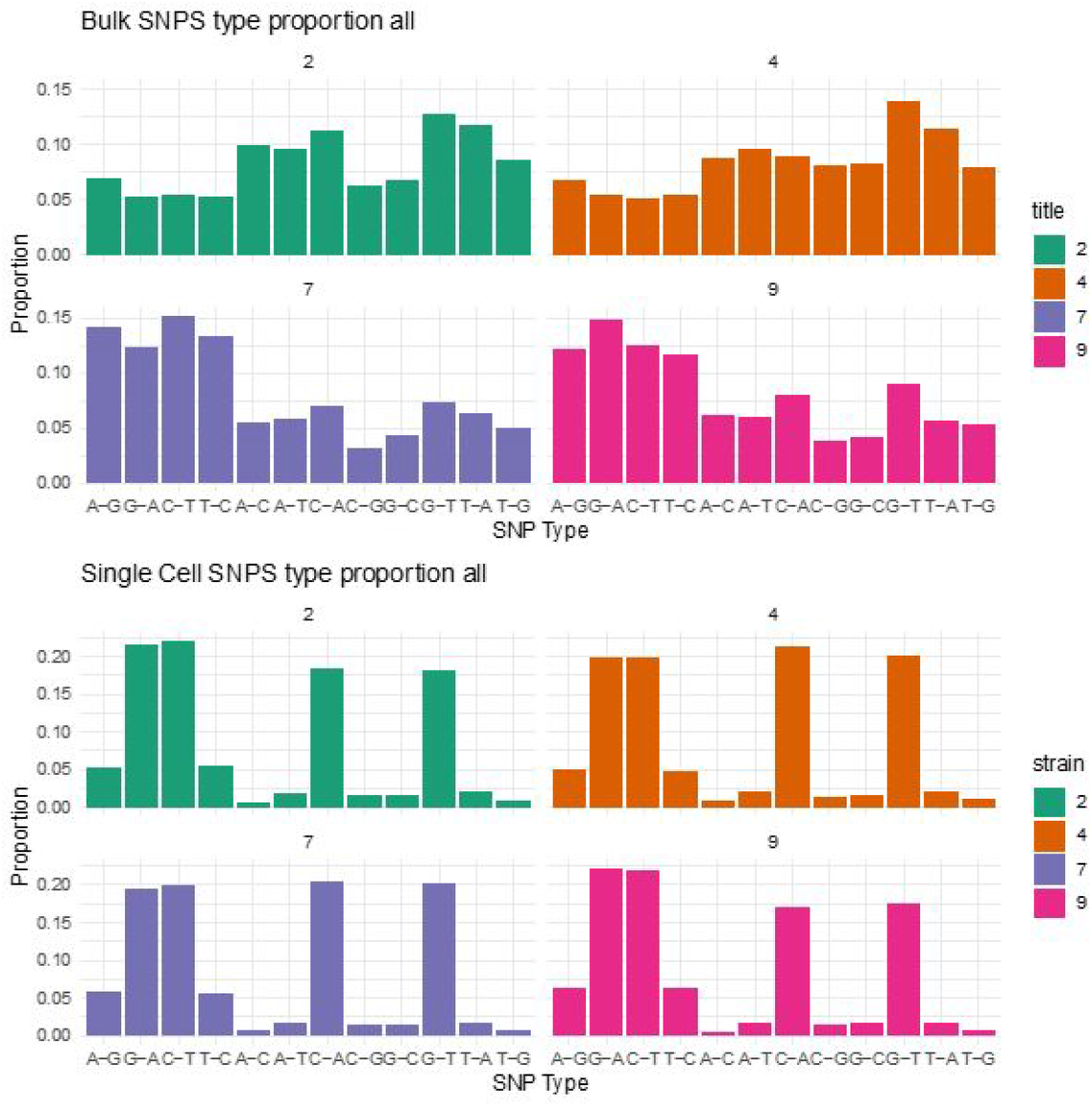
Proportion of SNP types seen in bulk (Top) and single cell sequencing (bottom) before filtering procedure. In the bulk sequencing data 2753 SNPS are called at 1500 unique genome sites for the four samples, SNP profiles show differences between the wild-type and hypermutator with more transitions are seen in the hypermutator (mean proportion = 0.53) than in the wild type (mean proportion = 0.23). In the single cell data 115286 SNPs are called at 99617 unique genome sites. SNP profiles are similar for all four samples and distinct to those seen in bulk sequencing. SNP types are dominated by G-A and C-T transitions (mean proportion 0.42) and C-A and G-T transversions (mean proportion 0.38)

**Supplementary Figure 3.**
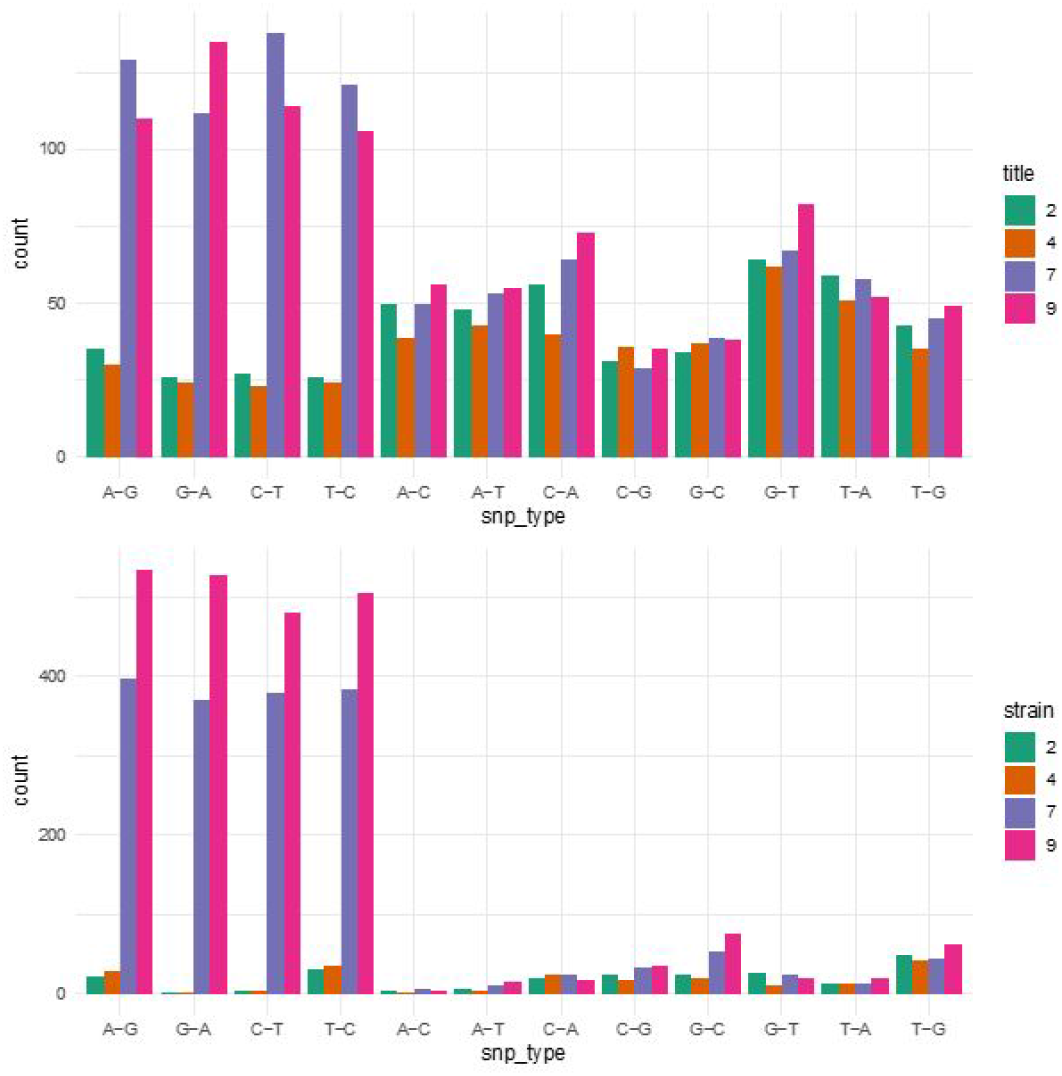
SNP profiles of bulk PCR free sequencing (top), and for filtered single cell SNPS derived from MDA (bottom). Single Cell SNPS were filtered by removing low frequency variants and G-A and C-T SNPs only seen in one single cell genome, resulting in 4448 SNPs at 1536 positions. Of these SNPs 234 (15%) of locations were also seen in bulk SNPs.

## Materials and Methods

### Bacterial cell culture and antibiotic stress

Initial ‘Barnyard’ experiments were performed using *S.* Enteritidis strain P125109 (Klemm et al., 2016) and *E. coli* strain MG1655 which constitutively expressed the yellow fluorescent protein YFP (Balaban et al., 2004). Two *S. enteritidis* strains were used in this study, PT4 P125109 [AM933172.1] (Klemm et al., 2016) and its isogenic *mutS* deletion mutant (included to potentially see more variation in a short period due to an elevated mutation rate). The MIC of ciprofloxacin was determined for both strains as being 0.08 mg/L. Both strains were repeatedly passaged in the presence and absence of 0.5 X the MIC of ciprofloxacin in LB broth, twice daily (am and pm) over two weeks. Parallel lineages were run with two dilution factors (1:1000 and 1:100000). The experiments completed between ~150 and ~300 generations (depending on the inocula). After each passage an aliquot (0.5ml) of each culture was stored with 15% glycerol and frozen to provide an archive of each time point for all lineages.

### AMR phenotyping

Susceptibility testing used the broth and agar dilution methods with Mueller-Hinton media following the guidelines and interpretation criteria provided by EUCAST (http://www.eucast.org/ast_of_bacteria/)

### Bulk DNA extraction

Cells were grown overnight at 37° C at 200 rpm in LB broth. The overnight culture was pelleted by centrifugation at 3,220 g for 15 minutes. The supernatant was discarded and the pellet was resuspended in 9mL of sterile PBS. Cells were lysed by adding 1mL of 20% SDS and 50 μL of a 20 mg/mL proteinase K solution before incubation at 37°C for 1 hour.

For DNA purification, 1 volume of phenol:chloroform:isoamyl alcohol mixture (25:24:1) was added to 1 volume of cell lysate and centrifuged at 3,220 g for 5 minutes. The aqueous (top) phase was recovered with a cut-end 1mL tip and transferred to a new clean tube. A volume of chloroform:isoamyl alcohol mixture (24:1) was added to the aqueous phase, and the tube was mixed gently by inverting until the content turned milky. The tube was centrifuged at 3,220 g for 5 minutes and the aqueous (top) phase was recovered once again with a 1mL cut-end tip and transferred to a new tube. To precipitate the DNA, 1/10 volumes of 3 M sodium acetate pH 5.2 and 2.5 volumes of ice-cold absolute ethanol were added to the tube before mixing gently by inverting until DNA precipitates. A Pasteur pipette hook was used to spool the DNA pellet and transfer it in a 2 mL tube containing 80% ethanol for a 5 minutes wash. The DNA pellet was then air dried for 5 minutes on the hook before being resuspended in 200 μL of ultrapure water, by incubation at room temperature overnight.

### Heterogeneity of susceptibility

To examine the heterogeneity of susceptibility within a population, 96 individual colonies were isolated from *mutS* lineages (exposed and control) and the MICs of ciprofloxacin and nalidixic acid were determined to identify the prevalence of resistance within the population. This confirmed selection of resistance in the exposed lineage, but also identified heterogeneity within the population with 13.6% of individual isolates being resistant (MIC >0.06mg/L) to ciprofloxacin in the exposed lineage (**Supplementary figure 1**), compared to none in the control. This data confirmed there was heterogeneity in the population and therefore the samples were a suitable test set for analysis by single cell sequencing.

### Bulk Illumina sequencing

Ten reference samples were sequenced on the Illumina NovaSeq platform with a PCR-free library preparation and generating 150 base paired end reads. Genomes were sequenced to an average depth of around 2700X.

### Reference genome

The chromosomal sequence of *Salmonella enterica* subsp. enterica serovar Enteritidis str. P125109 (AM933172) (Klemm et al., 2016) annotated using Prokka (version 1.11) (Seemann, 2014) was used as a reference genome.

### Variant Calling from Bulk Data

Read mapping to the reference and variant calling was performed using the breseq pipeline (Deatherage and Barrick, 2014), developed for the long-term *E. coli* evolution experiment. This pipeline uses bowtie2 (Langmead and Salzberg, 2012) for mapping and reports the fraction of reads that align to the reference genome. Polymorphisms present in at least one percent of reads with a read coverage of 10 on each strand at the variant position were called. Variants only seen on one strand were removed. The *breseq* pipeline also calculated various mapping and coverage statistics.

### Genome Coverage

The average total coverage was determined for all sequences from the *breseq* output. The total number of aligned bases was divided by the reference genome length. The evenness of coverage was determined by calculating the Lorenz curve for each sample https://github.com/yhoogstrate/bam-lorenz-coverage/tree/master/bin.

### Duplicate Reads

The number of duplicate reads per in the forward and reverse read sets was determined using dedupe.sh in bbmap-38.06.

### GC Content

The GC content of forward and reverse reads was determined using stats.sh in bbmap-38.06 (https://sourceforge.net/projects/bbmap/), the average of these values was then determined and used in subsequent analysis.

### Single bacterial isolation by FACS

Liquid cultures of bacterial cells were grown in Luria Bertani broth (LB) at 37°C and diluted in PBS. Bacterial cells were examined by size (FSC-A) and granularity (SSC-A) using Fluorescence-activated cell sorting (FACS, BD FACS Melody), and cells were then sorted into 96-well microplates with 100-cell and single-cell samples were included in our experiments. For the barnyard experiment, a mix of *Salmonella* and *E. coli* cultures was prepared and analysed by FACS, after each population was defined we proceeded to randomly sort cells of *E. coli, Salmonella* or a mix of both into individual wells of a 96-well plate.

### Whole Genome Amplification (WGA)

WGA of single cells was performed by multiple displacement amplification using the REPLI-g Single Cell kit (Qiagen, Valencia, CA) as per the manufacturer’s instructions but at one quarter of the recommended volumes. The amplified product was cleaned with AMPure Beads XP on a robotic platform (Biomek Nx, Beckman Coulter), quantified by Qubit (Thermo Scientific) and diluted to approximately 1 ng/μl before proceeding with library preparation.

### Single-cell sequencing library preparation

Sequencing libraries were prepared with Nextera XT kit (Illumina) as per the manufacturer’s instructions but at one quarter of the recommended volumes. The library product pooled and cleaned up using Ampure XP beads. The pooled library size was examined using a Bioanalyzer (Agilent) using the High Sensitivity DNA kit, and quantified by KAPA Library Quantification Kit (Roche).

### Single-cell genome sequencing and data analysis

For all single-cell experiments, initial paired-end (150 bp) sequencing was performed on the Illumina MiSeq platform (Nano flow cell) to assess library quality. For the barnyard experiment, no further sequencing was performed and read trimming and read alignment (BWA-MEM 0.7.17) were performed in Partek Flow (Partek Inc., St. Louis, USA). Sorted S. Enteritidis/mutS cells were subsequently sequenced using NovaSeq 6000 platform (SP flow cell, PE 150 bp reads).

### Single-cell variant calling

Read mapping to the reference and variant calling was performed using the breseq pipeline, as above. Polymorphisms present in at least one percent of reads with a read coverage of 10 on each strand at the variant position were called. Variants only seen on one strand were removed.

### Fixed SNP sites

The non-unique fixed SNP sites (SNPs present at a frequency greater than 0.9 and seen in more than one single cell genome) in the single-cell genomes were determined leading to 3452 SNPs at 357 unique locations and ultimately 316 SNP locations within genes. The alignment coverage at each of these positions was determined from the *breseq* bam files. Each SNP location could then be classified as SNP, reference or low-coverage for each single cell genome. For each culture (i.e. Hypermutator grown with ciprofloxacin), genomes with more than the mean plus one standard deviation positions out of the 316 candidate sites lacking sufficient coverage for SNP calling were excluded for phylogenetic reconstruction and population structure determination.

### Phylogenetic reconstruction from single-cell data

A sequence alignment of variant sites from above was used to generate a maximum likelihood phylogenetic tree, for each culture, with RAxML using the GTRCAT model implemented with an extended majority-rule consensus tree criterion (Stamatakis, 2006). RHierBaps (an R implementation of hierarchical Bayesian analysis of Population Structure) (Tonkin-Hill et al., 2018) was used to estimate population structure from the sequence alignment using two nested levels of molecular variation and up to 20 populations.

